# Annual Cycle Dampening and the Decrease Predictability of Water Level Fluctuations in a Dam-Regulated Neotropical Floodplain

**DOI:** 10.1101/824177

**Authors:** Jaques Everton Zanon

## Abstract

The flood pulse concept refers to seasonal variations in river water level and is the driving force in river-floodplain systems that ‘responsible for the existence, productivity and interactions’ of these system’s biota. This seasonal variation is inherent to river-floodplain systems and establishes a natural pattern of these ecosystems that has frequently been observed in nature. One particular river-floodplain system of interest is the Upper Parana River and its floodplain, whose upstream contains a reservoir cascade that caused profound alterations on its flooding regime by having diminished flood magnitude, but increased its frequency. In this study, I sought to explore the flood pulse condition in the Upper Paraná River Floodplain by using a set of state-of-the-art spectral and non-linear analyses and a time series of water level fluctuations (1968-2017) from this system. I divided the data into four periods: i) natural regime period, ii) transitional period, iii) dam cascade period, and iv) Primavera’s dam period. Spectral analysis demonstrated a decrease in the annual cycle amplitude, reflected in its power spectrum, which means a weakening in the difference between flood and drought events. Additionally, nonlinear dynamical analysis revealed a less deterministic and predicable behavior leading to more erratic fluctuations jeopardizing the temporal heterogeneity of that system.

## Introduction

Understanding the dynamics of water level fluctuations in floodplains is very important to elucidate the flood regime and its consequences (Lowe-McConnell, 1987). The flood pulse concept (Junk and others 1989) refers to seasonal variations in river water level and is the driving force in river-floodplain systems that are ‘responsible for the existence, productivity and interactions’ of these system’s biota. Consequently, the concept (flood pulse) attempts to relate the biota component with the environmental component of big river-floodplain systems. This seasonal variation is inherent to river-floodplain systems and establishes a natural pattern of these ecosystems that has frequently been observed in nature (Middleton 2002).

One particular river-floodplain system of interest is the Upper Parana River and its floodplain. That interest is because it represents the last stretch of this river that remains undammed in the Brazilian territory. It is located inside three Conservation Units (Ilha Grande National Park, Ivinheima State Park and Environmental Protection Area of ‘‘Ilhas e Várzeas do Rio Paraná’’) and since 1999, the Upper Paraná River and its floodplain have been included in the Long-Term Ecological Research (LTER) Program. However, upstream from this floodplain, is a reservoir cascade that caused profound alterations of the flooding regime, diminishing flood magnitude but increasing flood pulse frequency (Souza-Filho and others 2004). The alteration of the flood-pulse regime, reduces the seasonally flooded area and duration of floods leading to a change of systems dynamics (Ward and Stanford 1995; Souza-Filho 2009). The upper Paraná floodplain has its biochemical cycles intimately related to periodic increases of water level (Thomaz and others 1992).

One particular contribution to the understanding of damming history of this system was provided by Souza-Filho, (2009). The author categorized the time series of water level fluctuations into two periods: The period before 1972 is the Natural Regime Period and the period after 1972 is the Modified Regime Period (or Controlled Regime Period). The latter period is subdivided into three other intervals: the Transitional Period (from 1972 to 1981), the Regulated by the Cascade of Dams Effect Period (from 1982 to 1998) and the Regulated by the Porto Primavera Hydroelectric Power Plant Period (from 1999 to recent time).

Investigate the flood dynamics in each of these periods can provide insightful information about the modification caused by damming regulations. We can analyze these time series in the time domain using classical linear methods such as autocorrelation. Alternatively, they can be analyzed in the frequency domain, using spectral analysis. When analyzing a sample time series, analysis in both the time and frequency domains may be useful in highlighting different features of the data. That features can summarized using wavelet analysis. Some have recommended the use of the Fourier transform to decompose variation in ecological or environmental phenomena, such as hydrodynamics (Sabo and Post 2008). However, wavelets have the advantage over Fourier transforms in their scale independence and ability to examine multiple scales simultaneously (Torrence and Compo 1998). Most of the signals encountered in hydrology and earth sciences are characterized by intermittent or transient features that lead to nonstationary data series (Labat 2010). Due to their attractive properties, such as insensibility to nonstationary signals, wavelets have been explored for use in aquatic time series analysis being particularly useful for identifying the regularities (e.g. annual cycles) in a time series (Seuront and Strutton 2004; Labat 2010; Niu and Sivakumar 2013; Maity and others 2016).

Despite these known regularities characterized by annual flood cycles, there is an inherent nonlinear nature of hydrologic time series that comprehends an example of complex dynamic phenomenon that can jeopardize such regularities (seasonality) and prediction power (Sivakumar, 2009). This limitation makes classical linear approaches such as autocorrelations functions and classic time series tools not very appropriated to this type of data. In this case, the use of nonlinear time series analysis (NTSA) techniques is very important for attain information from this complex systems (Sivakumar and Berndtsson 2010; Huffaker and others 2017). The goal of any nonlinear dynamical analysis of a data series is to extract features of the dynamics of the underlying physical and chemical processes that produce that spatial pattern or time series. The basis of nonlinear analysis lies in powerful proposals put forward by Packard et al., (1980) and proved rigorously by Takens, (1981). The main task of nonlinear time series analysis is to extract information on the dynamical system from the observation of its evolution. This approach is different from the statistical one, in the sense that it can overcome typical limits of the traditional linear and statistical tools. As a result of this analysis one can estimate the level of determinism (and predictability) of the system and evaluate the amount of system’s order which can be used to gather information about the entropy of the system (Parrott 2004).

Since floodplain biotas are heavily influenced by flow regimes and their pristine flood events have been strongly modified by dams operations affecting their magnitude and predictability I intent in this study to analyze the flood pulse condition in the Upper Paraná River Floodplain. Using a set of state of the art spectral and non-linear analysis and a time series of water level fluctuations (from 1968-2017) from this system. I believe that the combination of those two approaches can give valuable information about the seasonality and predictability of flood in this system. I identified the dominant periodicities (e. g. annual cycles) and some non-linear features from each scenario proposed by Souza-Filho, (2009).

## Methods

### Study Area

I studied the water level fluctuations in the Upper Paraná River floodplain, one of the most dammed stretch of the Paraná River, suffering influences from several hydroelectric impoundments (Fig. 1). This floodplain is located at the border of the states Paraná state, Brazil and is part of a long-term ecological research project, which started in 2000 (see http://www.peld.uem.br/)

**Figure 1:**
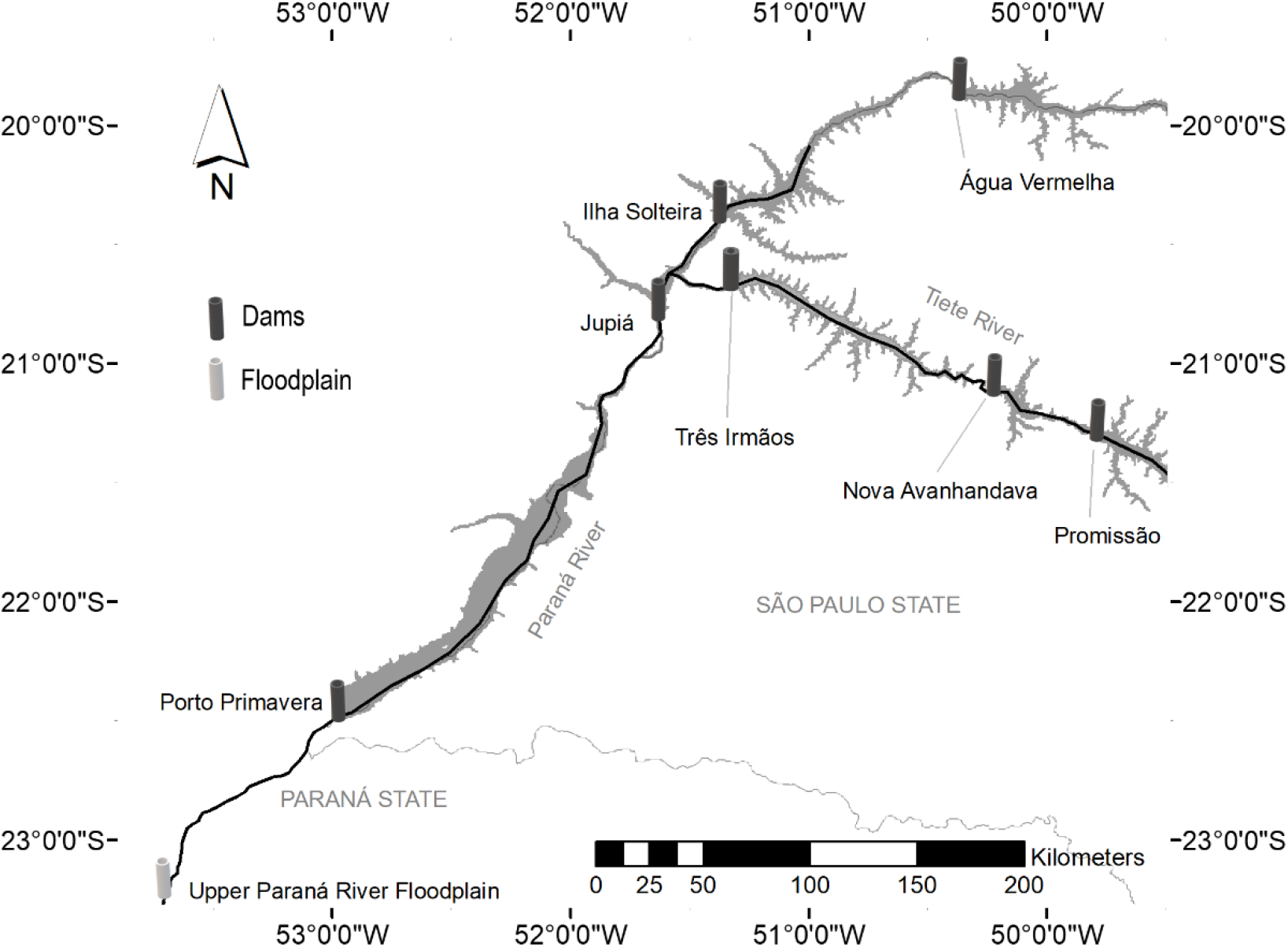
Upper Paraná River Floodplain location and the portrait of the damming cascade in the Paraná River, Brazil.

### Hydrologic time-series data

To initiate the study, I developed a hydrological database of river water levels fluctuations from the study area. This hydrologic data is public available and can be downloaded from the Agência Nacional de Águas (ANA; Brazil’s National Water Agency) using the Hydroweb platform querying for Porto São José hydrological station (www.ana.gov.br). The database includes daily measurements starting from January of 1964 to December of 2017 resulting in a 54 years of hydrological data. Subsequently, I categorize the database according to Souza-Filho, (2009): from 1964-72 is the Natural Regime Period; 1973-81 is the Transitional Period; 1982-98 is the Cascade of Dams Effect Period and 1999-recent time is the Primavera’s Dam period.

### Wavelet analysis

To access the different types of periodicity on the hydrological time series data, I used the wavelet transform method. In this analysis the original time series is compared to a mother wavelet function. There are plenty of mother wavelets (e.g. Mexican hat, Gaussian and Morlet). Particularly in this study, I used the Morlet wavelet due to its localization in both time and frequency and because it is widely use in climate data (SANTOS and others 2001; Grinsted and others 2004; Mwale and others 2004) and ecological data (Cazelles and others 2008). This type of analysis is a spectral technique that partitions the variability contained in a time series into variability associated with specific timescales. We used this method in each one of the four categories of time-series data with the intent to validate and compare the strength of periodicities in each period. For that, we used the WaveletComp package (Rosch and Schmidbauer 2014) in the R program (R Core Team 2019).

### Non-linear analysis

In order to study the evolution of water level fluctuations in each delimited period one can embed the water level time series to an N-dimensional vector space or Phase Space. This process recreates the dynamics of system conditions through the study of the revealed trajectories of the water level fluctuations in the Phase-Space. This is the cornerstone (thirst steps) to study dynamic systems. In this section, I will describe my approach, based on the concepts of the Nonlinear Time Series Analysis framework: the attractor reconstruction, the recurrence plot (RP), and the recurrence quantification analysis (RQA).

### Attractor reconstruction

The attractor of the water level underlying dynamics was reconstructed in phase space by the time delay vector method (Takens 1981; Abarbanel 1996). Starting from a time series [s_1_… s_N_], where s_i_ = s(i∆t) and ∆t is the sampling time, the system dynamics can be reconstructed using this method. The system’s trajectory is expressed as a new matrix **X** in which each row is a vector **x**_i_ = [s_i_, s_i+τ…_ s_i + (D_E−1_)τ_] of phase space and i = 1 … N - (D_E_−1) τ. The matrix **X** is characterized by two key parameters: the embedding dimension D_E_ and the delay time τ. The delay parameter τ can be estimated using the Mutual Average Information function (Thiel and others 2004) and the embedding dimension D_E_ by using False Nearest Neighbors (Abarbanel 1996). The False Nearest Neighbors is a dimensionality test that estimate the minimal number of dimensions while reducing the quantity of false nearest neighbors in limited number of dimensions. In other words, it estimates the number of state variables that are responsible for the system’s dynamics. This dimensionality test results in the minimum dimension at which the reconstructed attractor can be considered completely unfolded and there is no overlapping of the trajectories. If the dimensionality is to low, the attractor is not completely unfolded and the underlying dynamics cannot be investigated. This is the basis for transforming the one-dimensional time-series **s** into a multidimensional phase-space **X** (Fig. 2).

**Figure 2:**
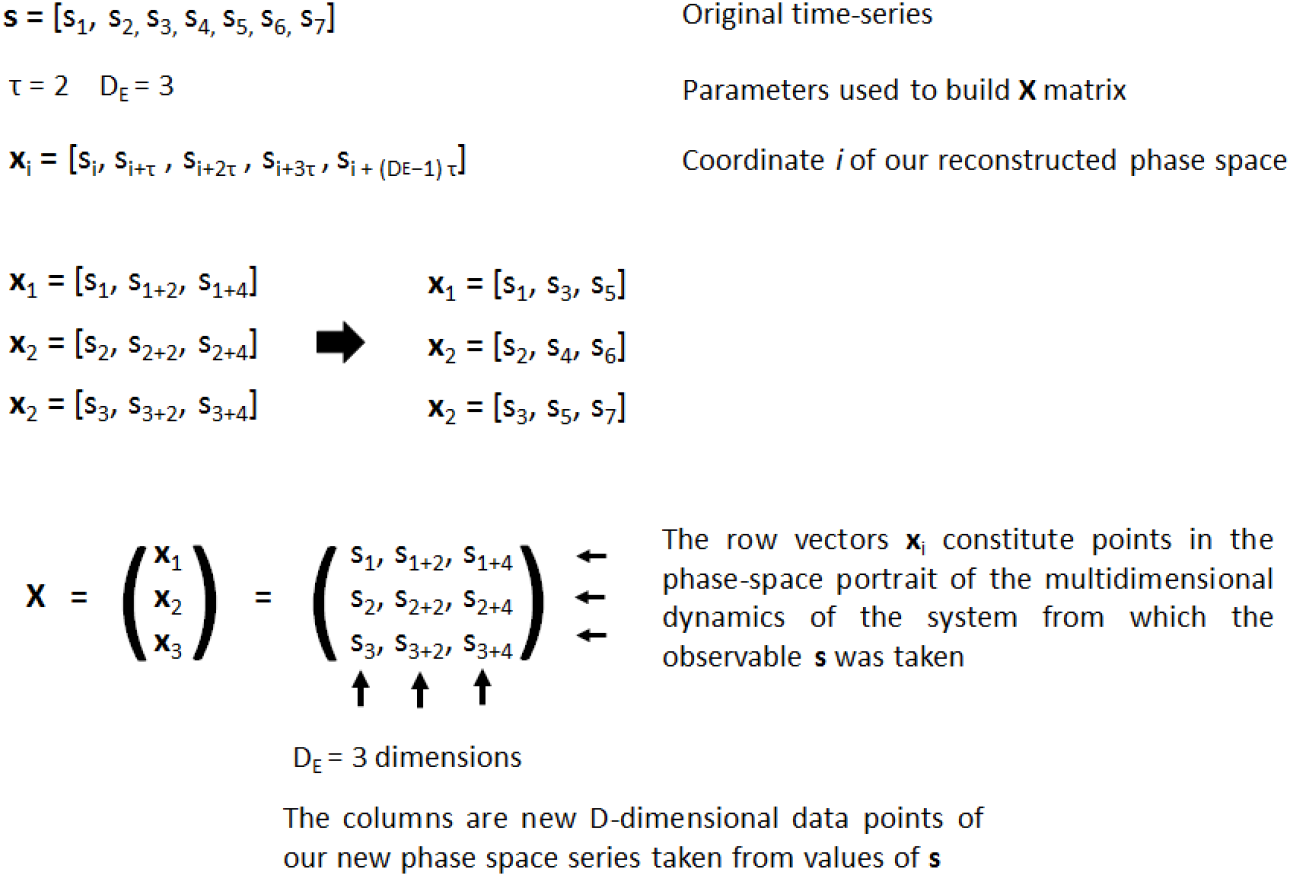
Phase space (attractor) reconstruction using Taken’s Theorem. Tau (**τ**) = time lag parameter and DE = number of dimensions needed to better represent the phase space)

### Time series analysis based on recurrence plots (RPs)

Once the matrix **X** is reconstructed based on the original time series, a common feature observed is the recurrence of its state vectors **x**_i_, it is important to clarify that in matrix **X** states with large temporal distances may be close in state space. To visualize such recurrences and proximities of states the recurrence plot (RP), proposed for the first time by Eckmann et al., (1987), is a visual tool able to identify temporal recurrences in multidimensional phase spaces. In the RP, any recurrence of state *i* with state *j* is pictured on a boolean matrix expressed by:

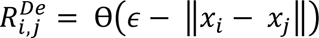

where xi, xj ∈ R^De^ are the embedded vectors, i, j ∈ N, Ɵ (⋅) is a step function and *ϵ* is an subjective threshold. Thus, in the resulting graph (see Fig.4) every nonzero entry of *R*_*i*,*j*_, thanks to the step function, is marked by a black dot in the position (i, j), otherwise, the plot are left blank. For instance, if the signal in one’s data is a sine wave (a complete deterministic and periodic pattern), then recurrences will occur with a period equal to that of the signal; the recurrence plot comprises continuous diagonal lines that are continuous thought the plot. The lack of any diagonal lines suggest that the signal is stochastic (se Fig. 1 in Hobbs and Ord 2018).

### Recurrence quantification analysis (RQA)

The RP provides a visual manner for inspecting the recurrent behavior of variables and various complex patterns, which is very useful for time series analysis as they provide a graphical impression of the dynamics of the study system (Marwan and others 2007). However, beyond superficial impressions of stochasticity and determinism, the plots provide no quantification of such processes. Recurrence quantification analysis (RQA) is based on deriving quantitative measures from the distribution of lines on the recurrence plot. It is a method initially develop in theoretical physics (Eckmann and others 1987) and applied to various non-linear problems numerous and diverse fields of study (Webber and others 2009). I calculated seven RQA quantitative descriptors: the percent determinism, the Shannon entropy (ENT), the mean length of the diagonal lines (Lmean), the maximum diagonal line length (Lmax), the trend (TREND), laminarity (LAM) and trapping time (TT). For a detailed discussion of these measures, please see Webber and Zbilut (2005) and Marwan and others (2007) however in Tab. 1 I provided some information.

**Table 1:**
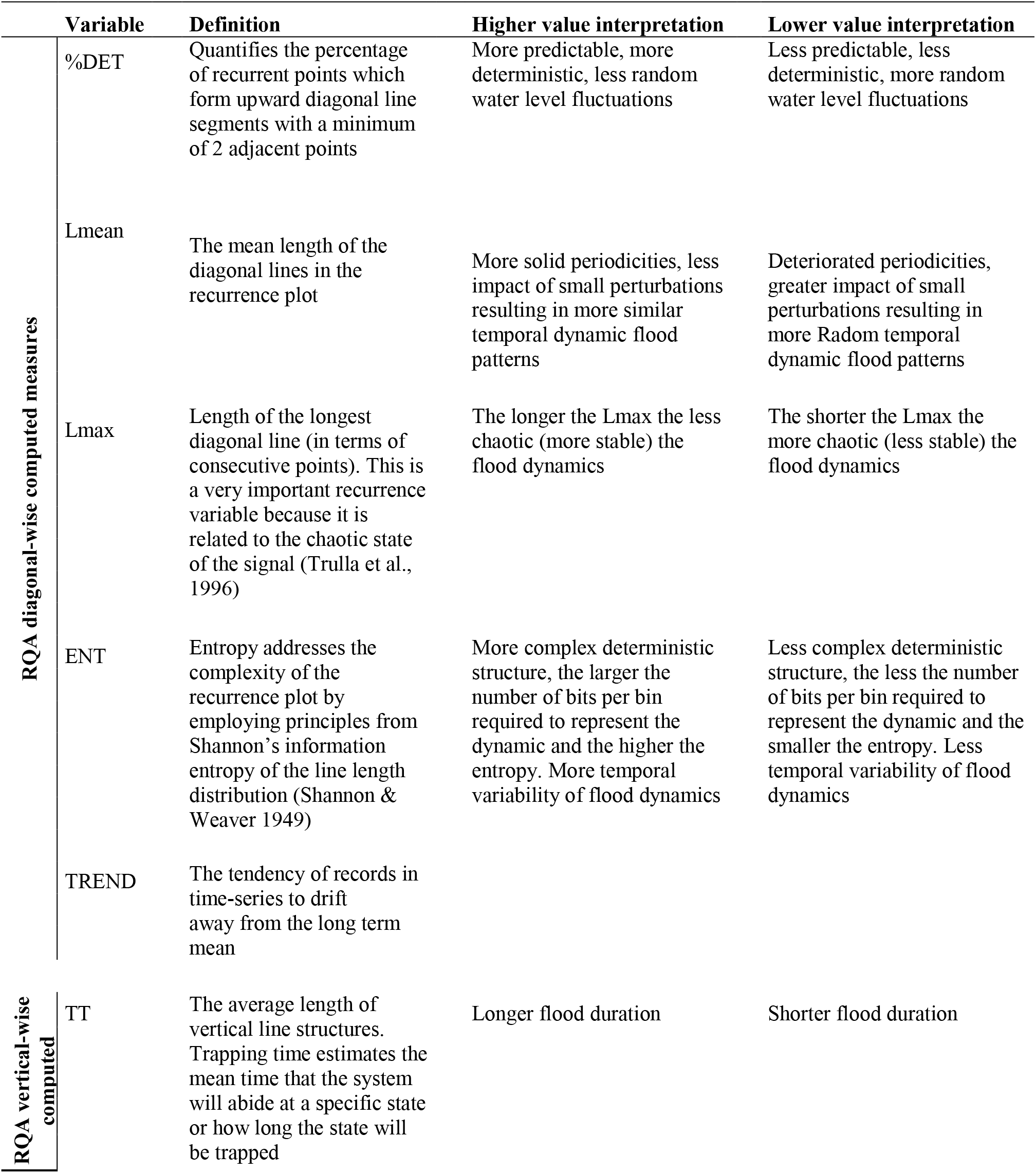
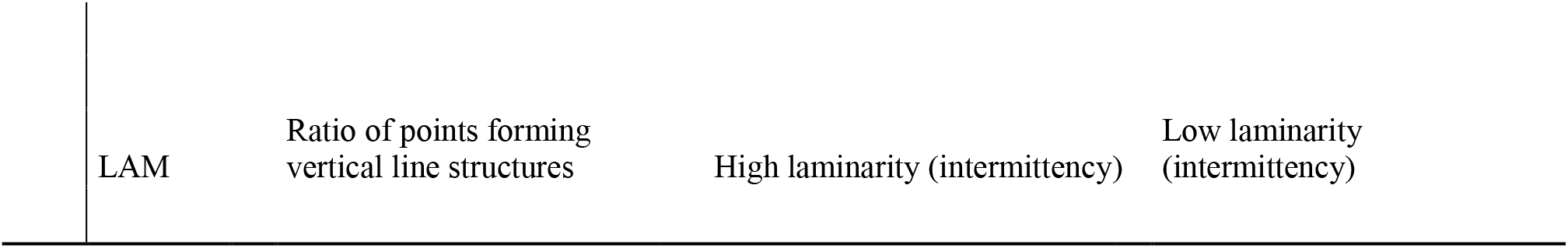
Dynamics invariants used in the Recurrence quantification Analysis for each period in the time-series.

|  | Variable | Definition | Higher value interpretation | Lower value interpretation |
| --- | --- | --- | --- | --- |
| RQA diagonal-wise computed measures | %DET | Quantifies the percentage of recurrent points which form upward diagonal line segments with a minimum of 2 adjacent points | More predictable, more deterministic, less random water level fluctuations | Less predictable, less deterministic, more random water level fluctuations |
|  | Lmean | The mean length of the diagonal lines in the recurrence plot | More solid periodicities, less impact of small perturbations resulting in more similar temporal dynamic flood patterns | Deteriorated periodicities, greater impact of small perturbations resulting in more Radom temporal dynamic flood patterns |
|  | Lmax | Length of the longest diagonal line (in terms of consecutive points). This is a very important recurrence variable because it is related to the chaotic state of the signal (Trulla et al., 1996) | The longer the Lmax the less chaotic (more stable) the flood dynamics | The shorter the Lmax the more chaotic (less stable) the flood dynamics |
|  | ENT | Entropy addresses the complexity of the recurrence plot by employing principles from Shannon's information entropy of the line length distribution (Shannon & Weaver 1949) | More complex deterministic structure, the larger the number of bits per bin required to represent the dynamic and the higher the entropy. More temporal variability of flood dynamics | Less complex deterministic structure, the less the number of bits per bin required to represent the dynamic and the smaller the entropy. Less temporal variability of flood dynamics |
|  | TREND | The tendency of records in time-series to drift away from the long term mean |  |  |
| RQA vertical-wise computed | TT | The average length of vertical line structures. Trapping time estimates the mean time that the system will abide at a specific state or how long the state will be trapped | Longer flood duration | Shorter flood duration |

|  |  |  |  |
| --- | --- | --- | --- |
| LAM | Ratio of points forming vertical line structures | High laminarity (intermittency) | Low laminarity (intermittency) |

## Results

The visual inspection of the seasonality of the data on each period can demonstrate key differences since the start of the impoundments (Figure 3). Visually there is an equivalence of water level values towards the end of the time series (2017). This lack of difference within the year period can be an indication to seasonality weakening of the flood pulse. The attractor reconstruction revealed high dimensionality (D_E_) in the four periods (Table 2). The period with higher dimensionality was the period mainly influenced by the dam cascade in which six state variables are responsible for its attractor. The remaining periods all were composed by 5 state variables. It is worth to recall that each state variable is a different dimension in phase space (Fig. 2) and can be recognized as a controlling variable not sampled or unknown in the system.

**Table 2:** Embeddology parameters used in phase space (attractor) reconstruction. Dimensionality was estimated using Mutual Average Information procedure, time lag was estimated using False Nearest Neighbors procedure and the Radius was chosen in order to maintain the recurrence rate (RR) at 10%.

| Period | Dimensionality ($D_E$) | Time lag ($\tau$) | Radius ($\epsilon$) | Time Series Length (obs.) |
| --- | --- | --- | --- | --- |
| Natural | 5 | 110 | 160 | 3288 |
| Transitional | 5 | 60 | 125 | 3653 |
| Dam cascade | 6 | 100 | 100 | 6572 |
| Primavera | 5 | 140 | 80 | 6993 |

**Figure 3:**
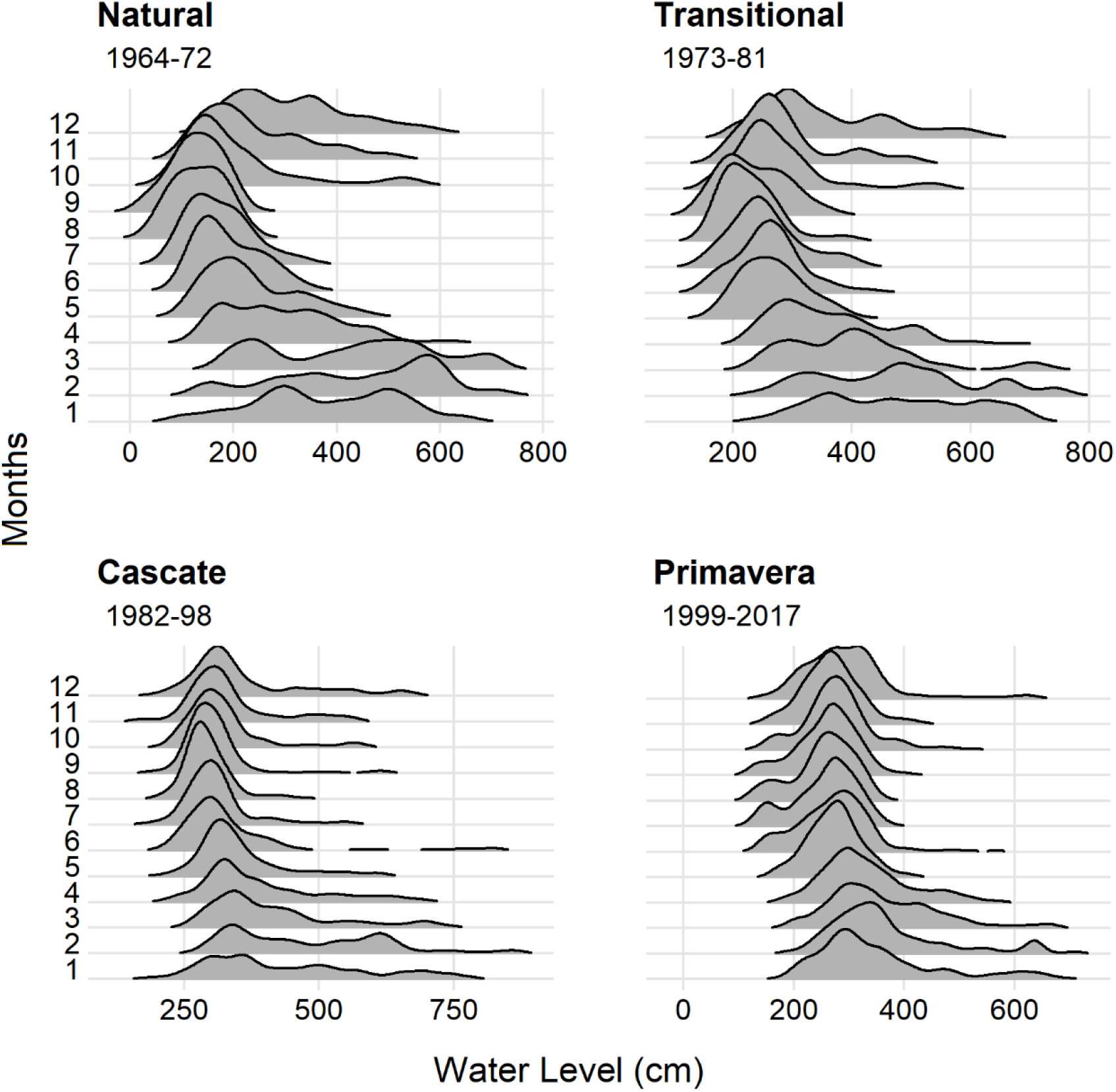
Density graphs of intra-annual water level values (cm) in the four periods of the historical time-series.

### Raw water level time series (Figure 4a)

The original time series of the selected time periods are displayed in Fig 4a. Visually, it is virtually impossible to infer anything about the system dynamics from these figures, however, some visual features can be highlighted. In the Natural period, some clearer periods are visible with more periodic peaks. This pattern seems to dissolve in the next periods where strong peaks are followed by epochs with noisier structures. So, this gives the impression that the damming operations reflects in noisier and possible chaotic dynamics. Therefore, wavelet analysis and RQA which allows the analysis of periodicities and chaotic behavior was applied to these datasets in order to have insights into the dynamics of the flood pulse.

**Figure 4:**
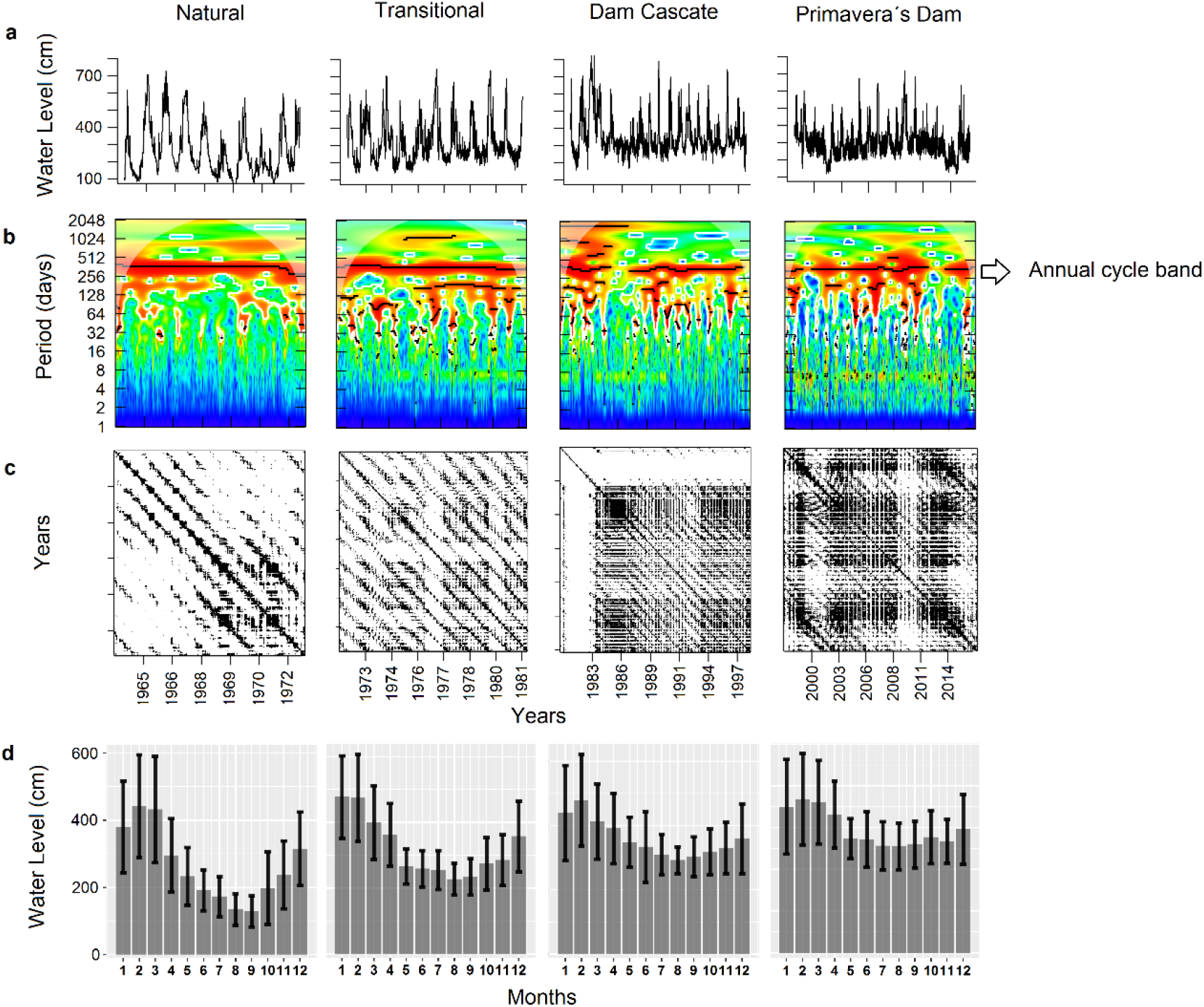
Raw time series and results from spectral (wavelets) and nonlinear analysis (RP). a) Raw water level time series; b) Wavelet graphics; c) Recurrent Plots (RPs) and d) Bar plots with standard deviation.

### Wavelet analysis (Figure 4b)

Clearly there is a predominance of the yearly water level cycle (the flood pulse *per se*) in each of the four periods (red band across the entire four wavelets graphs). The dominant band in concentrated in the 370 days period across all four periods, however, as can be noted in Fig. 5, its absolute power level has decreased at each studied period. This reflects what in observed in Fig. 3 the decrease of the seasonality strength. One interesting thing that can be observed in the wavelets portrait in the appearance of numerus sub annual bands in the Transitional, Dam cascade and Primavera periods. This indicates the presence of (added to the annual cycle) semiannual, quarterly, monthly and even weekly cycles (in Primavera graph one can note small spots in the 7-8 days period band). This demonstrates strong changes in the water level periodicities in the floodplain system.

**Figure 5:**
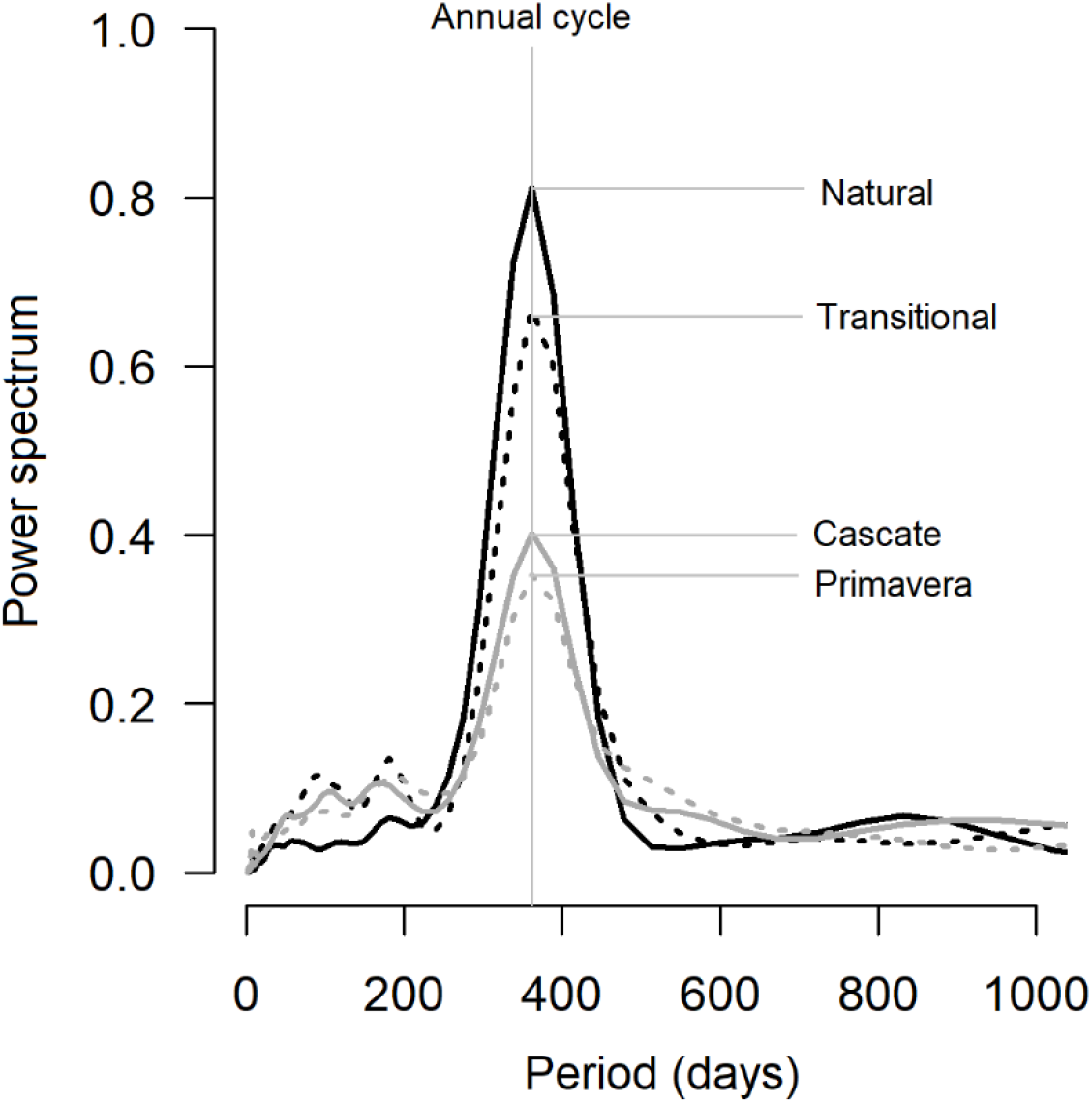
Wavelet power spectrum from each time period studied. All the periods had a power peak ate 369 days (approx. a year period = annual cycle).

### Recurrence Plots and Recurrence Quantification Analysis (Figure 4c)

In addition, I compared patterns of RPs of the four periods. The dynamic characteristics of the daily water level series cam be differentiated by comparing their RPs, in which the recurrence patters are critically different form each other. Long diagonal lines in the first period (Natural) was more common than the other periods, suggesting higher predictability in the Natural period. In fact, this period presented higher determinism and Lmean values. A decrease of diagonal lines cam be observed in both RP and RQA and the RP exhibited noisier signals in the subsequent time periods. In figure 4c in the dam cascade RP a white band can be noted. This indicate that the systems states in this time interval are rare or far from normal signifying a strong nonstationarity in water level dynamics. All dynamic invariants (except TREND) decreased towards more recent time periods (Figure 6).

**Figure 6:**
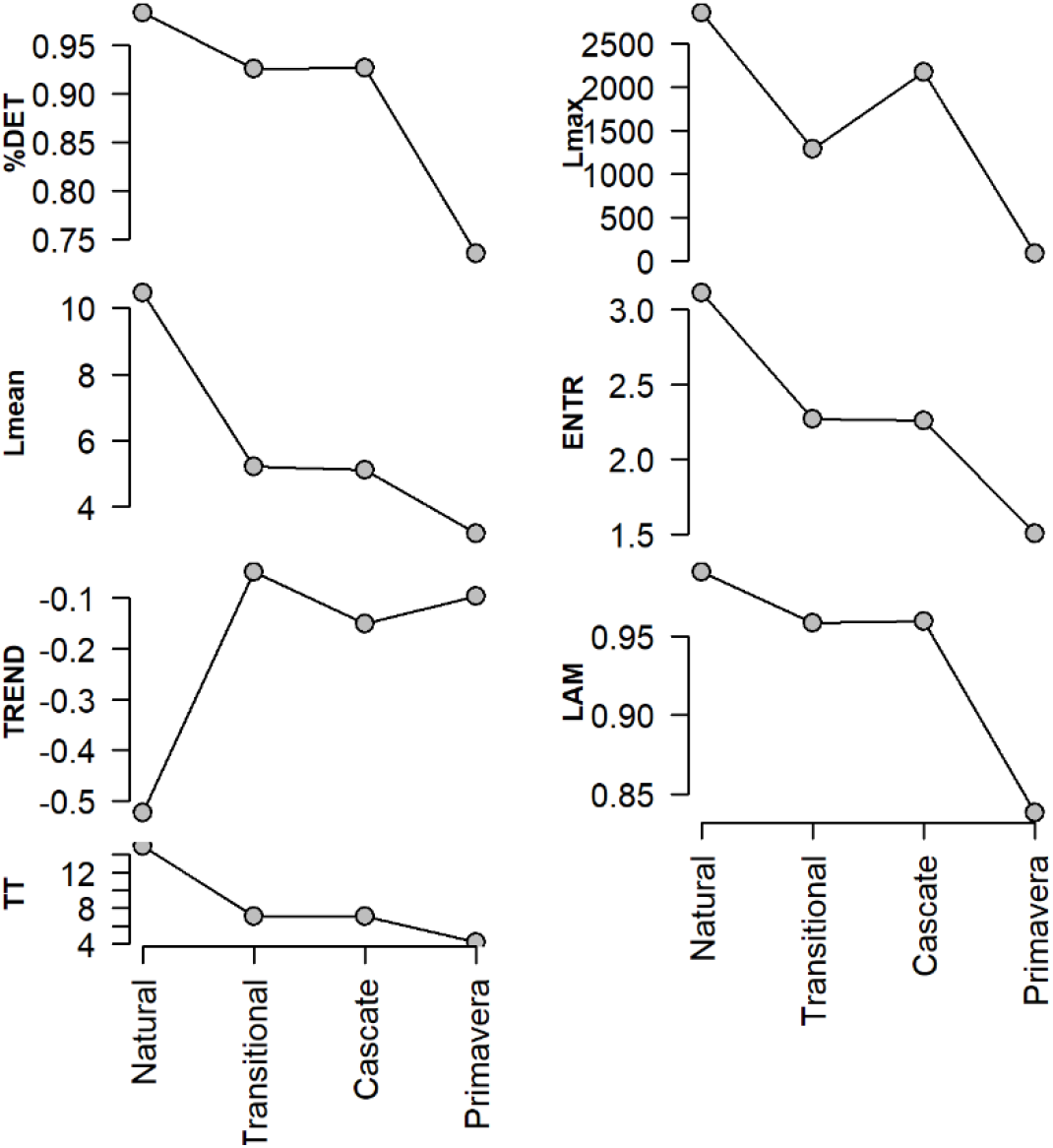
Dynamic invariants resulted from Recurrence Quantification Analysis (RQA) of Recurrence Plots presented in Fig. 4. (For description and interpretation of these variables please see Tab. 1)

## Discussion

Water levels are one of the key categories of stress affecting aquatic ecosystems in general (Baron and others 2002) and in the Paraná floodplain in particular (Algarte and others 2009; Padial and others 2009; Gomes and others 2011; Dunck and others 2013, 2016; de Assis Murillo and others 2019). Despite being an already well explored subject, the flood information is important to ecological structure and function, it is important that we asses a better understanding of the fluctuating water levels portrait (Allan and others 2013). With the help of wavelet and RQA we accessed quantitative results for several variables, which allowed a comprehensive description of the non-linear behavior of the flood pulse in the floodplain system. This was useful for comparisons across all four periods, not only with respect to the annual cycle problem but also studying deterministic behavior, transient processes, and trends in those time series. This allowed us to gain key information not available thought classical procedures and will assuredly help us to better conservations and management practices of the system at hand.

### Flood Pulse Spectra

One interesting aspect addressed by this study is the periodicity of the flood pulse. Periodicity is a fundamental pattern in many natural time series and, the periodicity of water level fluctuation of the Parana River Floodplain suffered critical changes across the studied time periods. Despite observing an appreciable power in the spectrum of time-frequency analysis (wavelet) obtained form all four time periods, it has been weakened across the years (Fig. 5). This mean that the seasonal hydrological regime is losing its power, in other words, the floods and the droughts are leveling (Fig. 3) causing a hydrologic stability in the floodplain.

This type of change is directly linked to the provision of ecosystem services, and the consequences of such leveling are worrisome. For instance, Nielsen and others (2013) demonstrated experimentally a critical reduction in biotic diversity as a consequence of flood smoothing. Fewer plant species, reduction of overall abundance, and simplifications of ecological communities are some of the critical effects described by the authors. Other authors indicate that WLFs have the potential to influence sediment–water nutrient exchange, which may influence system productivity (Steinman and others 2014). Similarly, the downstream riverine ecosystem health are also affected by flood stability. For instance, Resende and others (2019) detected significant loses of Amazonian floodplain forests as a consequence of intense flood regulation promoted by damming the Uatumã river.

### Diagonal-wise dynamic invariants (DET, Lmax, ENT and TREND)

According to common stochastic hydrology theory (Yevjevich 1972, 1974, 1987; Sivakumar 2017), both deterministic and random components are included in hydrologic time series. Deterministic components, mainly including periodic process and transient process (trends), are generated by certain deterministic physical mechanisms. The usual way of looking for order in a time series is to compute its power spectrum (Degn and others 1987). Generally, in a pure random process, the power spectrum fluctuates erratically around a constant value, stipulating that are no prominent frequency explaining any more of the variance of the sequence than any other frequency. So, we can identified such periodic deterministic process using wavelet spectral analysis (Smith and others 1998; Labat 2008). In natural conditions, the flood pulse can be perfectly predicable, and characterized by a monomodal flood pulse with an annual cycle of high and low water periods (Junk and others 1989). Determinism and order are prevalent in such natural annual flood pulse. The decrease of such determinism can point out a shift to a more stochastic structure in flood dynamics. Although we observed frequency predominance (annual flood pulse) in all studied periods (Fig. 5) a less deterministic structure was present towards the end of the time-series (Fig. 6).

As a measure of flood instability, a lower DET value indicates less predictability of water level fluctuations (see Tab. 1). DET is the percentage of total recurrences that form diagonal lines in the recurrence plot and the mean diagonal line (Lmean) are both related to predictability in a positive way (Webber Jr. and others 2005). Similarly to DET Lman also decreased and reflected such positive link between then. Such results indicates a noisier or a more ‘random-looking’ structure in our data. The time series periods was separated accordingly to dam operations. Such damming generates a well know cascade effect impacting the entire river-floodplain system (Agostinho and others 2004). The combine cascading of events can strongly amplify the impact caused by single events in terms of extension of the affected area and damage level. The final cascade impact can be shattering and need to be prudently evaluated in terms of planning and management (Zuccaro and others 2018).

The diagonal lines in the RP correspond to parallel trajectory sections, so the extent of these lines is somehow related to the divergence behavior of the dynamical system. Lmax corresponds to the length of the longest diagonal line segment in the plot and short diagonal lines is an indicative of system’s divergence (i.e. mathematical chaos) (Eckmann and others 1987; Trulla and others 1996). In a simple way, chaos refers to disorder or the absence of some kind of order. A chaotic system is a complex, aperiodic (no system recurrences) and sensitive to its initial conditions and implies strong restrictions on the ability to predict future states. In that sense, the power spectrum provide convincing distinction between periodic and aperiodic systems. Therefore, the study system is a periodic system and maintain its periodicity in the four study periods (Fig. 5), however, it has lost a great deal of its power spectrum as already discussed in previous section. While it would need more specific analysis to detect the presence of chaos in our time-series data (Ellner and Turchin 1995; Marwan 2010; Sivakumar 2018) it is arguable that as clearly reflected by the values of the dynamic invariants (Fig. 6) the amount of disorder (noise) in the flood dynamics has increased towards the end of the time-series periods.

Entropy (ENT) is a measure of disorder, it quantifies the ‘amount of uncertainty’ in the time-series (Shannon 1948) and it is a quantity of complexity. Higher values of entropy in a Recurrence Plot corresponds to a more uniform distribution of lengths of diagonal lines (Fig. 4c ‘*Natural’*) resulting determinist structures of greatly variable lengths. On the other hand, low entropy values results from a distribution of diagonal lines’ lengths focused around a few persistent values (less variable values – Fig. 4c *Primavera*). This few persistent values can be related to a certain homogeneity in the water level distribution values towards the last study period culminating in more homogenous density distribution (fig. 3) and Recurrent Plots (Fig. 4c). The results indicates more homogeneus point’s distribution towards the last study period. Stationary time-series provides somewhat homogeneous recurrence plots, while nonstationary signals causes major changes in points distribution of recurrence points resulting in bright visible areas in the plot (Marwan and Kurths 2002). This information is also captured in the TREND variable. The lesser the TREND the more stationary the system. This indicates that the water levels fluctuations in the PIARP is getting more stationary jeopardizing its flood temporal variability (Tonkin and others 2017).

### Vertical-wise dynamic invariants (%LAM and TT)

Both %LAM and TT indexes decreased towards the last study period. They are related to flood intermittency (periodicity). Intermittent behavior reflects the water level fluctuation that exhibits changes in dynamics from fluctuating (slow changes; 1^st^ period - Natural) to relatively stationary (fast changes; last period - Primavera). This results are complementary to the spectral and RQA diagonal-wise invariants since it also show a less periodic (more stationary) behavior.

In the present study, a statistical framework based on spectral, nonlinear theory and recurrence analysis of dynamical systems has been proposed in order to quantitatively identify the temporal characteristics of flood pulses in UPRF. The statistics support more stochastic, nonlinear and unpredictable structure in the flood pulse dynamics. The analysis presented helps to statistically discern the temporal patterns of the flood pulse phenomena and characterize the transitions observed. Moreover, the above mentioned characteristics of flood pulse dynamics may have significant implications to future impoundments underling an extraordinary situation of shifts between droughts and floods conditions. Such anthropogenic stressors are altering the natural periodicity of abiotic and biotic events. Dams are the present example example of anthropogenic stressors that severely altered seasonality in the study system. This altered seasonality can be harmful, jeopardizing the ability of organisms to harmonize their cycles with the seasonal phenomena (Lytle and Poff 2004; Wernberg and others 2013) and critically reducing the temporal diversity (Tonkin and others 2017). Yet, infrequent and unpredictable floods can act as a disturbance alternating the system productivity (Parsons and others 2005). The Upper Paraná River floodplain is indispensable for Brazilian biodiversity, because this region shelters an important fraction of the original biota of the basin. Three protected areas are present in this region, indicating their relevance for biodiversity conservation; however, their ecological integrity is threatened by a series of upstream reservoirs and an inappropriate use of the watershed.

## References

Abarbanel HDI. 1996. Analysis of Observed Chaotic Data. Springer http://link.springer.com/10.1007/978-1-4612-0763-4

Agostinho AA, Thomaz SM, Gomes LC. 2004. Threats for biodiversity in the floodplain of the Upper Paraná River : effects of hydrological regulation by dams. Ecohydrol Hydrobiol 4:255–6.

Algarte VM, Siqueira NS, Murakami EA, Rodrigues L. 2009. Effects of hydrological regime and connectivity on the interannual variation in taxonomic similarity of periphytic algae. Braz J Biol 69:609–16. http://www.ncbi.nlm.nih.gov/pubmed/19738967

Allan JD, McIntyre PB, Smith SDP, Halpern BS, Boyer GL, Buchsbaum A, Burton GA, Campbell LM, Chadderton WL, Ciborowski JJH, Doran PJ, Eder T, Infante DM, Johnson LB, Joseph CA, Marino AL, Prusevich A, Read JG, Rose JB, Rutherford ES, Sowa SP, Steinman AD. 2013. Joint analysis of stressors and ecosystem services to enhance restoration effectiveness. Proc Natl Acad Sci 110:372–7.

de Assis Murillo R, Corrêa Alves D, dos Santos Machado R, Silveira MJ, Fidanza Rodrigues K, Thomaz SM. 2019. Responses of two macrophytes of the genus Polygonum to water level fluctuations and interspecific competition. Aquat Bot.

Baron JS, Poff NL, Angermeier PL, Dahm CN, Peter H, Hairston NG, Jackson RB, Johnston CA, Richter BD, Steinman AD, Hairston NG, Jackson RB, Johnston CA. 2002. Meeting Ecological and Societal Needs for Freshwater Stable URL : http://www.jstor.org/stable/3099968 Linked references are available on JSTOR for this article : MEETING ECOLOGICAL AND SOCIETAL NEEDS FOR FRESHWATER. 12:1247–60.

Cazelles B, Chavez M, Berteaux D, Ménard F, Vik JO, Jenouvrier S, Stenseth NC. 2008. Wavelet analysis of ecological time series. Oecologia 156:287–304. http://www.ncbi.nlm.nih.gov/pubmed/18322705. Last accessed 21/07/2014

Degn H, Holden A V., Olsen LF. 1987. Chaos in biological systems. (Degn H, Holden A V., Olsen LF, editors.). Boston, MA: Springer US http://link.springer.com/10.1007/978-1-4757-9631-5

Dunck B, Algarte VM, Cianciaruso MV, Rodrigues L. 2016. Functional diversity and trait-environment relationships of periphytic algae in subtropical floodplain lakes. Ecol Indic 67:257–66. http://dx.doi.org/10.1016/j.ecolind.2016.02.060

Dunck B, Bortolini JC, Rodrigues L, Rodrigues LC, Jati S, Train S. 2013. Functional diversity and adaptative strategies of planktonic and periphytic algae in isolated tropical floodplain lake. Brazilian J Bot 36:257–66. http://link.springer.com/10.1007/s40415-013-0029-y. Last accessed 02/09/2014

Eckmann J., Kamphorst SO, Ruelle D. 1987. Recurrence Plots of Dynamical Systems. Europhys Lett 4:973–7. http://stacks.iop.org/0295-5075/4/i=9/a=004?key=crossref.09fbeb6883f90a0adb050fbd7323bcd5

Ellner S, Turchin P. 1995. Chaos in a Noisy World: New Methods and Evidence from Time-Series Analysis. Am Nat 145:343–75. https://www.journals.uchicago.edu/doi/10.1086/285744

Gomes LC, Bulla CK, Agostinho a. a., Vasconcelos LP, Miranda LE. 2011. Fish assemblage dynamics in a Neotropical floodplain relative to aquatic macrophytes and the homogenizing effect of a flood pulse. Hydrobiologia 685:97–107. http://link.springer.com/10.1007/s10750-011-0870-6. Last accessed 13/06/2013

Grinsted A, Moore JC, Jevrejeva S. 2004. Application of the cross wavelet transform and wavelet coherence to geophysical time series. Nonlinear Process Geophys 11:561–6. http://www.nonlin-processes-geophys.net/11/561/2004/

Hobbs B, Ord A. 2018. Nonlinear dynamical analysis of GNSS data: quantification, precursors and synchronisation. Prog Earth Planet Sci 5.

Huffaker R, Bittelli M, Rosa R. 2017. Nonlinear Time Series Analysis With R. Oxford University Press

Junk WJ, Bayley PB, Sparks RR. 1989. The flood pulse concept in river-floodplain systems. In: Dodge DP, editor. Proceedings of the International Large River Symposium (LARS), Canadian Journal of Fisheries and Aquatic Sciences. NRC research press. pp 110–27. http://swrcb2.swrcb.ca.gov/waterrights/water_issues/programs/bay_delta/bay_delta_plan/water_quality_control_planning/docs/sjrf_spprtinfo/junk_et_al_1989.pdf

Labat D. 2008. Wavelet analysis of the annual discharge records of the world’s largest rivers. Adv Water Resour 31:109–17.

Labat D. 2010. Wavelet analyses in hydrology. In: Sivakumar B, Berndtsson R, editors. Advances in Data-Based Approaches for Hydrologic Modeling and Forecasting. World Scientific Publishing Company. pp 371–410.

Lytle DA, Poff NLR. 2004. Adaptation to natural flow regimes. Trends Ecol Evol 19:94–100.

Maity R, Suman M, Verma NK. 2016. Drought prediction using a wavelet based approach to model the temporal consequences of different types of droughts. J Hydrol 539:417–28. http://dx.doi.org/10.1016/j.jhydrol.2016.05.042

Marwan N. 2010. How to avoid potential pitfalls in recurrence plot based data analysis. 21:1003–17. http://arxiv.org/abs/1007.2215 http://dx.doi.org/10.1142/S0218127411029008

Marwan N, Carmen Romano M, Thiel M, Kurths J. 2007. Recurrence plots for the analysis of complex systems. Phys Rep 438:237–329.

Marwan N, Kurths J. 2002. Nonlinear analysis of bivariate data with cross recurrence plots. Phys Lett A 302:299–307. https://linkinghub.elsevier.com/retrieve/pii/S0375960102011702

Middleton BA. 2002. Flood pulsing in wetlands: restoring the natural hydrological balance. John Wiley & Sons

Mwale D, Gan TY, Shen SSP. 2004. A new analysis of variability and predictability of seasonal rainfall of central southern Africa for 1950-94. Int J Climatol 24:1509–30.

Nielsen DL, Podnar K, Watts RJ, Wilson a. L, Watts KPRJ. 2013. Empirical evidence linking increased hydrologic stability with decreased biotic diversity within wetlands. Hydrobiologia 708:81–96. http://link.springer.com/10.1007/s10750-011-0989-5. Last accessed 16/04/2013

Niu J, Sivakumar B. 2013. Scale-dependent synthetic streamflow generation using a continuous wavelet transform. J Hydrol 496:71–8. http://dx.doi.org/10.1016/j.jhydrol.2013.05.025

Packard NH, Crutchfield JP, Farmer JD, Shaw RS. 1980. Geometry from a Time Series. Phys Rev Lett 45:712–6. https://link.aps.org/doi/10.1103/PhysRevLett.45.712

Padial AAA, Carvalho P, Thomaz SM, Boschilia SM, Rodrigues RB, Kobayashi JT, Carvalho ÆP, Becker R, Josilaine RÆ, Kobayashi T, Nin E. 2009. The role of an extreme flood disturbance on macrophyte assemblages in a Neotropical floodplain. Aquat Sci 71:389–98. http://link.springer.com/10.1007/s00027-009-0109-z. Last accessed 28/03/2013

Parrott L. 2004. Analysis of simulated long-term ecosystem dynamics using visual recurrence analysis. Ecol Complex 1:111–25.

Parsons M, McLoughlin CA, Kotschy KA, Rogers KH, Rountree MW. 2005. The effects of extreme floods on the biophysical heterogeneity of river landscapes. Front Ecol Environ 3:487–94.

Proulx R, Côté P, Parrott L. 2009. Multivariate recurrence plots for visualizing and quantifying the dynamics of spatially extended ecosystems. Ecol Complex 6:37–47.

R Core Team. 2019. R: A Language and Environment for Statistical Computing. http://www.r-project.org

Resende AF de, Schöngart J, Streher AS, Ferreira-Ferreira J, Piedade MTF, Silva TSF. 2019. Massive tree mortality from flood pulse disturbances in Amazonian floodplain forests: The collateral effects of hydropower production. Sci Total Environ 659:587–98. https://doi.org/10.1016/j.scitotenv.2018.12.208

Rosch A, Schmidbauer H. 2014. WaveletComp : A guided tour through the R-package.:1–38.

Sabo JL, Post DM. 2008. QUANTIFYING PERIODIC, STOCHASTIC, AND CATASTROPHIC ENVIRONMENTAL VARIATION. Ecol Monogr 78:19–40. http://doi.wiley.com/10.1890/07-1861.1

Santos CAG, Galvão C de O, Suzuki K, Trigo RM. 2001. MATSUYAMA CITY RAINFALL DATA ANALYSIS USING WAVELET TRANSFORM. Proc Hydraul Eng 45:211–6. http://joi.jlc.jst.go.jp/JST.Journalarchive/prohe1990/45.211?from=CrossRef

Seuront L, Strutton PG. 2004. Scaling Methods in Aquatic Ecology.

Shannon C. 1948. A Mathematical Theory of Communication. Bell Syst Tech J 27:379–423.

Sivakumar B. 2009. Nonlinear dynamics and chaos in hydrologic systems: Latest developments and a look forward. Stoch Environ Res Risk Assess 23:1027–36.

Sivakumar B. 2017. Chaos in Hydrology. Dordrecht: Springer Netherlands http://link.springer.com/10.1007/978-90-481-2552-4

Sivakumar B. 2018. Chaos Identification and Prediction Methods. In: Sivakumar B, editor. Chaos in Hydrology. Dordrecht: Springer Netherlands. pp 173–98.

Sivakumar B, Berndtsson R. 2010. Advances in Data-Based Approaches for Hydrologic Modeling and Forecasting. World Scientific Publishing Company http://ebooks.worldscinet.com/ISBN/9789814307987/9789814307987.html

Smith LC, Turcotte DL, Isacks BL. 1998. Stream flow characterization and feature detection using a discrete wavelet transform. Hydrol Process 12:233–49.

Souza-Filho EE. 2009. Evaluation of the Upper Paraná River discharge controlled by reservoirs. Brazilian J Biol 69:707–16. http://www.ncbi.nlm.nih.gov/pubmed/19738976. Last accessed 14/10/2013

Souza-Filho EE, Rocha PC, Comunello E, Stevaux JC. 2004. Effects of the Porto Primavera Dam on Physical environment of the downstream floodplain. In: Thomaz SM, Agostinho AA, Hahn NA, editors. The Upper Parana River and its Floodplain: Physical Aspects, Ecology and Conservation. Leiden: Backhuys Publishers. pp 55–74.

Steinman AD, Ogdahl ME, Weinert M, Uzarski DG. 2014. Influence of water-level fluctuation duration and magnitude on sediment–water nutrient exchange in coastal wetlands. Aquat Ecol 48:143–59. http://link.springer.com/10.1007/s10452-014-9472-5. Last accessed 10/09/2014

Takens F. 1981. Detecting strange attractors in turbulence. pp 366–81. http://link.springer.com/10.1007/BFb0091924

Thiel M, Romano MC, Read PL, Kurths J. 2004. Estimation of dynamical invariants without embedding by recurrence plots. Chaos 14:234–43.

Thomaz SM, Lansac-Tôha FA, Roberto M do C, Esteves FA, Lima AF. 1992. Seasonal variation of some limnological factors of lagoa do Guarana : a varzea lake of the High Rio Parana, state of Mato Grosso do Sul, Brazil - 38924.pdf. Brazil Rev Hydrobiol 25:269–76. http://horizon.documentation.ird.fr/exl-doc/pleins_textes/cahiers/hydrob-trop/38924.pdf

Tonkin JD, Bogan MT, Bonada N, Rios-Touma B, Lytle DA. 2017. Seasonality and predictability shape temporal species diversity. Ecology 98:1201–16.

Torrence C, Compo GP. 1998. A Practical Guide to Wavelet Analysis. Bull Am Meteorol Soc 79:61–78.

Trulla LL, Giuliani A, Zbilut JP, Webber CL. 1996. Recurrence quantification analysis of the logistic equation with transients. Phys Lett A 223:255–60. http://linkinghub.elsevier.com/retrieve/pii/S0375960196007414

Ward J V., Stanford JA. 1995. Ecological connectivity in alluvial river ecosystems and its disruption by flow regulation. Regul Rivers Res Manag 11:105–19. http://doi.wiley.com/10.1002/rrr.3450110109

Webber CL, Marwan N, Facchini A, Giuliani A. 2009. Simpler methods do it better: Success of Recurrence Quantification Analysis as a general purpose data analysis tool. Phys Lett Sect A Gen At Solid State Phys 373:3753–6. http://dx.doi.org/10.1016/j.physleta.2009.08.052

Webber CL, Zbilut JP. 2005. Recurrence quantification analysis of nonlinear dynamical systems. In: Riley MA, Van Orden G, editors. Tutorials in contemporary nonlinear methods for the behavioural sciences. pp 26–92.

Webber Jr. CL, Zbilut JPP, Webber Jr CL. 2005. Recurrence quantification analysis of nonlinear dynamical systems. Tutorials Contemp nonlinear methods Behav Sci:26–94. http://www.nsf.gov/sbe/bcs/pac/nmbs/chap2.pdf

Wernberg T, Smale DA, Tuya F, Thomsen MS, Langlois TJ, De Bettignies T, Bennett S, Rousseaux CS. 2013. An extreme climatic event alters marine ecosystem structure in a global biodiversity hotspot. Nat Clim Chang 3:78–82. http://dx.doi.org/10.1038/nclimate1627

Yevjevich V. 1972. Stochastic Process in Hydrology. Water Resources Publications

Yevjevich V. 1974. Determinism and Stochasticity in Hydrology. J Hydrol 22:225–38.

Yevjevich V. 1987. Stochastic models in hydrology. Stoch Hydrol Hydraul 1:17–36.

Zuccaro G, De Gregorio D, Leone MF. 2018. Theoretical model for cascading effects analyses. Int J Disaster Risk Reduct 30:199–215.

